# An experimental manipulation of species’ phenologies overturns competitive hierarchies

**DOI:** 10.1101/573204

**Authors:** Christopher Blackford, Rachel M. Germain, Benjamin Gilbert

## Abstract

Ecological theory produces opposing predictions about whether differences in the timing of life history transitions, or ‘phenology’, promote or limit coexistence. Phenological separation is predicted to create temporal niche differences, increasing coexistence, yet phenological separation may competitively favour one species, increasing fitness differences and hindering coexistence. We experimentally manipulated relative germination timing, a critical phenological event, of two annual grass species, *Vulpia microstachys* and *V. octoflora*, to test these contrasting predictions. We parameterized a competition model to estimate within-season niche differences, fitness differences, and coexistence, and to estimate coexistence when among-year fluctuations of germination timing occur. Increasing germination separation caused parallel changes in niche and fitness differences, with the net effect of weakening within-year coexistence. Both species experienced a competitive advantage by germinating earlier, strongly enough to allow the generally inferior competitor to exclude the other with at least a four day head start. The overall consequence of germination separation was to limit coexistence within a given year, although among-year variation in relative timing of germination was sufficient to support long-term coexistence. Our results clarify how phenological differences structure competitive interactions, and highlight the need to quantify among-year variation in these differences to better understand species coexistence.

## Introduction

Species in many ecological communities show striking differences in the seasonal phenology of life history events, but the consequences of phenological differences for species coexistence are widely debated (Rabinowitz et al. 1981). Classic models predict that differences in phenology lead to reduced niche overlap among species, promoting coexistence (Gotelli and Graves 1996; Albrecht and Gotelli 2001). However, earlier phenology may also reduce resources available to later individuals or lead to size-structured competitive asymmetries, reducing the possibility of coexistence (Godoy and Levine 2014). Resolving these conflicting hypotheses is essential in this era of global change—species’ phenologies are shifting with climate change at different rates (Edwards and Richardson 2004; Scranton and Amarasekare 2017; Kharouba et al. 2018) and it is unclear how the fitness consequences of those shifts will play out in competitive environments (Yang and Rudolf 2010).

Coexistence theory offers a conceptual framework to understand how competitive interactions change when species differ in phenology. Specifically, Chesson (Chesson 2000) proposed two types of competitive differences that have opposing effects on coexistence, and can be quantified and then associated with specific traits (Kraft et al. 2015), such as phenology (Godoy and Levine 2014): ‘niche differences’ and ‘fitness differences’. Niche differences are present when intraspecific competition exceeds interspecific competition, thus introducing negative frequency-dependence that prevents any one species from dominating a community, stabilizing coexistence. By contrast, fitness differences are competitive asymmetries that give one species an advantage over the other and thus act to preclude coexistence. The combined effects of niche differences and fitness differences determine whether each species in a competitive pair can increase from low density when the other is abundant, and thus whether coexistence or exclusion are predicted (Fig. 1). Thus, the effects of phenological differences between competing species on coexistence is quantifiable by how they contribute to niche differences, fitness differences, or both.

**Figure 1.**
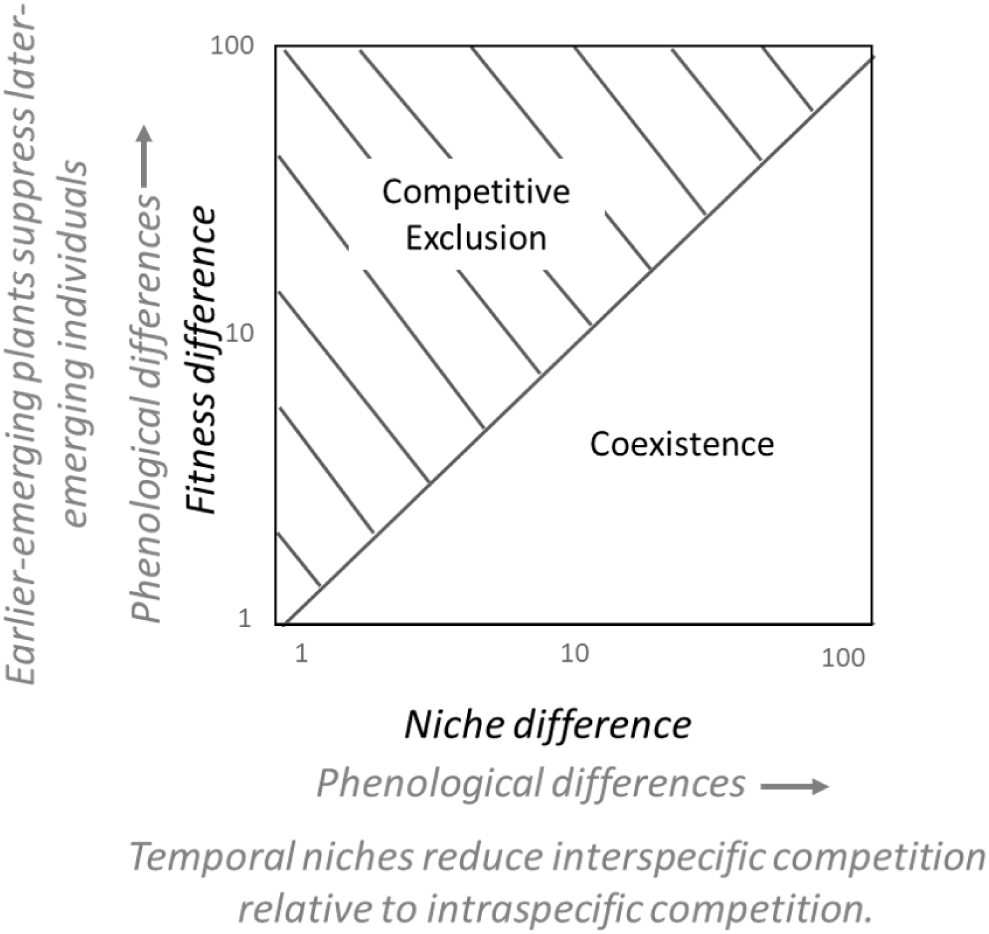
Opposing hypotheses about the effect of phenological differences on coexistence. Conceptual models on the effect of phenological differences (grey) mapped onto a coexistence framework (Adler et al. 2007). Coexistence occurs below the 1:1 line, when niche differences are high and fitness differences are low; note that although niche differences are typically presented as 1-*ρ*, causing the coexistence threshold to be non-linear, our presentation of 1/*ρ* places niche and fitness differences on the same scale, thus linearizing this threshold.

Research has provided mixed support for predictions of increased and decreased coexistence with phenological differences. Using annual plant communities from California, Godoy and Levine (2014) contrasted niche and fitness differences among plant species that differ in timing of flowering: early, mid, or late in the growing season. They found that, although phenological differences did increase niche differences, they contributed most strongly to fitness differences in favour of late-flowering plants (Godoy and Levine 2014). This study provides some of the clearest evidence about how phenological differences among co-occurring species map onto their competitive differences. However, the phenological differences observed were correlated to a suite of competitive traits, and the authors suggest these correlated traits (rooting depth and biomass) may ultimately be responsible for the observed effect of phenology (Godoy and Levine 2014). Additionally, they considered broad differences in phenology, such as summer vs. winter life histories, a magnitude of difference among species that is unlikely to shift with climate change or even interannual variation in climate. An experimental approach that manipulates phenology directly is necessary to isolate its effects, especially as shifting climatic regimes alter phenological responses independently of other traits

(Kharouba et al. 2018).Germination is a key phenological event for plants, as its timing determines if seedlings will grow in a tolerable climate and sets the competitive arena each plant will face (Donohue 2003), affecting fitness (Akiyama and ågren 2013). Not surprisingly, germination responds plastically to climate (Young et al. 2001; Levine et al. 2011), suchthat fluctuating climatic conditions can alter the absolute and relative timing of germination among species (Young et al. 1981). The composition of the neighbouring seed community also influences germination timing (Goldberg et al. 2001; Lortie and Turkington 2002). For example, Dyer et al. (2008) observed that a native California bunchgrass, *Nassella pulchra*, altered its timing of germination when seeds of competing species were present, with approximately half of all species causing *N. pulchra* germination to accelerate. This separation of germination timing might be an adaptive response to avoid interspecific competition (Young et al. 2017), especially given that competition tends to be particularly high during the emergence life stage (Goldberg et al. 2001; Chu and Adler 2015). If this hypothesis is correct, species should show greater niche differences and lower fitness differences when their germination is temporally segregated from other species. However, the alternate possibility, that early germination generates competitive differences that benefit earlier species, has also been observed in several studies (Harper et al. 1961; D’Antonio et al. 2001; Abraham et al. 2009; Grman and Suding 2010), although it is unclear if such advantages of early germination are symmetric or benefit some species more than others. Most studies of fine-scale variation in germination phenology do not test their net effects on niche differences, fitness differences, or coexistence, limiting their inferences (e.g. Young et al. 2001).

In this study, we experimentally isolate the effects of differences in germination timing on the coexistence of a congeneric pair of Mediterranean annual grasses, *Vulpia microstachys* ((Nutt.) Munro) and *V. octoflora* ((Walter) Rydb.). Germination phenology offers a unique opportunity for experimental manipulation as it is straightforward to induce for many species, and in doing so, clearly separates germination phenology from other traits. We manipulated the relative germination timing of these two *Vulpia* species, allowing each to germinate up to ten days in advance of the other and used an additive competition design to parameterize an annual plant model (Godoy and Levine 2014; Germain et al. 2016). We then determined (1) the effect of differences in germination timing on niche differences, fitness differences, and coexistence, and (2) how these outcomes change if differences in germination timing fluctuate from year to year. We show that early germination generally confers a competitive advantage, and when germination phenology varies among years, for example, due to climatic variability, the influence of phenology on coexistence differs on short (within year) and long (among year) timescales.

## Methods

### Study species and competition experiment

*Vulpia microstachys* and *V. octoflora* are generalist grasses that are widely distributed in California and commonly co-occur (Brooks 2000; Hoste 2013). As winter annuals, they germinate in late fall under cool, wet conditions and complete their life cycle by mid-summer. *Vulpia microstachys* germinates faster than *V. octoflora* under identical environmental conditions (Fig. S1), making it plausible that differences in their germination schedules promote coexistence. *Vulpia microstachys* has been shown to be competitively dominant to *V. octoflora* due to a lower resource requirement (R*) for several limiting resources (HilleRisLambers et al. 2010) and possibly due to its larger seeds. However, both species frequently persist together in mixed communities (HilleRisLambers et al. 2010).

Seeds of *V. microstachys* and *V. octoflora* were germinated in separate petri dishes over three weeks. Each petri dish contained 30 seeds of one species (seed density of 0.47 seeds/cm^2^) on filter paper, moistened with a 0.15% (v/v) solution of Previcur fungicide to suppress fungal growth that could interfere with germination. There were 130 replicate Petri dishes per species, which were sealed and kept under greenhouse conditions simulating a Californian winter. Daytime temperatures were set to maintain a 20ºC/15 ºC day/night temperature schedule with a 10 hour day length provided by supplemental high intensity discharge (HID) lighting. These conditions were maintained throughout plant growth.

When seedlings emerged (i.e., the moment the radicle broke through the seed coat), they were transplanted to 0.65 L cone-shaped pots (6.9 cm diameter, 25.4 cm depth) of sandy-loam soil, with the radicle slightly buried. We manipulated phenology by planting germinants into the same pot on different days. Pilot studies suggested that *V. octoflora* reaches full germination approximately 5 days after *V. microstachys* (Fig. S1) so to mimic realist germination differences, we constructed five treatments where *V. microstachys* would be planted −10, −5, 0, 5, or 10 days after of *V. octoflora*. To control for any effect of taking early/late-germinating individuals and to ensure enough seeds had germinated to conduct our experiment, seedlings were transplanted when total germination of each species reached 50%, which our pilot studies predicted would occur within 1-2 days for *V. microstachys* and 5-6 days for *V. octoflora* (Fig. S1). Because of natural variation in the exact time 50% germination was achieved, some of our germinants in this study were not planted precisely at 5 days intervals but instead were planted within a 24 hour window of that period. Given that this temporal window is small relative to our treatments, we do not account for this variation in our analysis and going forward refer to our treatments as −10, −5, 0, 5, and 10 days. There were 20 replicate pots of each treatment (20 pots5 treatments = 100 total pots), with pots containing 11 seedlings of each species to produce an overall density of 22 seedlings per pot, or 0.59 seedlings/cm^2^.

We also planted three monoculture control treatments per species (six treatments total) with 11 seedlings per pot; this additive design allowed us to parameterize our competition model (described below). There were ten replicate pots per monoculture treatment planted concurrent to the two-species competition (experiment day 0, 5, and 10) to correspond to the competition experiment, totalling 60 pots (10 pots ×3 planting dates × 2 species = 60 monoculture pots total). The monoculture treatment allowed us to estimate species’ population growth rates in the absence of interspecific competition, and also to quantify any changes in plant growth caused by the absolute date of planting rather than relative timing.

Growing conditions in the greenhouse were set according to earlier studies which included these same species, facilitating model parameterization (Germain and Gilbert 2014; Germain et al. 2016, 2018b). Following planting, 80 mL of water was added to each pot every three days via a drip irrigation system, and 75 mL of 1500 ppm 20-20-20 NPK fertilizer was added after four weeks but prior to flowering. Separately for each species, all mature seed produced in each pot was collected, counted, and weighed. We then used these data to calculate finite rates of increase (number of viable seeds produced per plant) and mean mass per seed (mg/seed). Germination tests were conducted to assess the proportion of seeds that germinated (parameter *g*_*i*_ in eqn. 1, details below). Although competition models assume the effects of competition will manifest through the number of seeds produced, competition might also impact the mass of individual seeds (Germain et al. 2018a); we also measured mass per seed to test this possibility. The experiment lasted until all plants had produced seed and senesced.

### Determining niche and fitness differences

Our experiment was designed to parameterize a Beverton-Holt annual plant competition model, which has previously been shown to capture the competitive dynamics of these species (Germain et al. 2016), and then used this model to estimate niche and fitness differences (Godoy and Levine 2014). We first conducted tests to determine which population parameters were influenced by germination separation. To do so, we tested the effects of germination separation on species’ finite rates of increase (number of viable seeds produced per plant), maximum percent germination of seeds, and seed mass. For these tests, we used generalized linear mixed effects models (‘lme4’ package in R) with species, germination separation, and their interaction included as fixed effects. Experimental ‘pot’ was included as a random factor to account for the lack of independence of measurements performed on both species in a single pot. Germination separation was treated as a continuous variable of the number of days that *V. microstachys* germinated relative to *V. octoflora* (with treatments replicated at −10, −5, 0, 5, 10 days separation).

The Beverton-Holt model that we fit is:

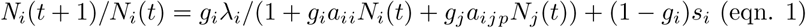

where the finite rate of increase of species *i, N*_*i*_(*t* +1)/*N*_*i*_(*t)*, is a function of the maximum rate of increase in the absence of competition (λ_*i*_), germination rate (*g*_*i*_), survival rate of ungerminated seeds (*s*_*i*_), the per capita intraspecific competitive impact (α*_ii_*), and the per capita interspecific competitive effects of species *j* on species *i* (α*_ijp_*). The subscript *p* denotes the value of α*_ij_* specific to a particular phenology (germination) treatment. The results of our linear models revealed no effect of planting date on seed production in the monoculture treatments (see Results and Fig. S2) justifying parameterizing our competition model with interspecific competition (α*_ij_*, α*_ji_*), but not intraspecific competition (α*_ii_*, α*_jj_*) or maximum rate of increase (λ), changing as a function of germination separation. Because the Beverton-Holt model is symmetric, dynamics of species *j* are represented by switching subscripts *i* and *j* in eqn. 1.

We estimated parameters all six parameters (three λ and α parameters per species) by fitting eqn. 1 to our data using a Bayesian approach, assuming each was lognormally distributed. All α parameters were constrained to be positive and uninformative priors were used (for all log(α), mean = 0, precision = 0.01, where precision is 1/variance). The λ priors for each species were taken from a study designed to estimate maximum rates of increase for these species in the same growing conditions as the current experiment, with mean log(λ) = 5.4 and 7.0 and precision = 4.7 and 4.0 for *V. microstachys* and *V. octoflora* respectively (Germain and Gilbert 2014). Parameter *g* was estimated from our germination trial data, and for our multi-year simulations (described below) we estimated *g* experimentally by conducting subsequent germination tests on the seeds produced in our experiment. Because measuring the survival rate of ungerminated seeds (*s*) in a greenhouse setting was not possible, we repeated all analyses with two extreme values (*s* =0 and *s*=1) to determine the sensitivity of our results to this parameter. For within-year calculations (eqns. 2-4), this parameter did not qualitatively change our results, so we only report *s*=0. The Bayesian model allowed us to estimate credible intervals on our hyperparameters (the composite parameters that jointly determine niche differences, fitness differences, and invasion growth rates; described below). A maximum likelihood approach can also be used estimate parameters and hyperparameters (described in the Supplementary Materials). This second approach gave similar point estimates but does not have the ability to generate confidence intervals so we report only the Bayesian results.

Once we parameterized eqn. 1, we used the parameter estimates to calculate niche differences, fitness differences, and to predict coexistence as their net effect (Fig. 1; Godoy and Levine 2014). When all viable seeds germinate, niche differences (1/ρ) in the Beverton-Holt model is:

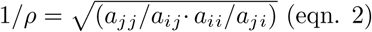

And the pairwise fitness difference (*κ*) is given as:

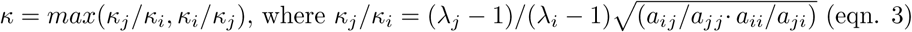

Coexistence is predicted to occur when *κ* < 1/*ρ* (Godoy and Levine 2014; Germain et al. 2016), meaning that fitness differences are less than niche differences. When this condition (*κ* < 1/*ρ*) is met, populations of each species are expected to increase when at low density and the other is at its equilibrium abundance. We note that niche differences are often presented as 1-*ρ*, but we use the definition in eqn. 2 allow a clearer graphical interpretation of coexistence outcomes by putting niche and fitness differences on the same scale (i.e., a linear coexistence threshold in Fig. 1); whenever the point defined by eqn. 2 exceeds that defined by eqn. 3, both species are expected to increase when initially at low densities. When some viable seeds fail to germinate, eqn. 3 changes such that the first fraction becomes (*η*_*j*_ − 1)*/*(*η*_*i*_−1), where *η*_*j*_ = (*λ*_*j*_*g*_*j*_)*/*(1 − *s*_*j*_ + *s*_*j*_*g*_*j*_) (Godoy and Levine 2014). The high levels of germination in our study (97% and 86% for *V. microstachys* and *V. octoflora* respectively) caused *ηλ*. The coexistence criteria *κ* < 1/*ρ* specifies the conditions necessary for both species to have positive population growth rates when at low density. Population growth rates under these conditions are referred to as ‘invader growth rates’ (Siepielski and McPeek 2010), which we can also solve directly using the equation:

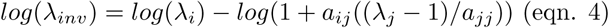

When both species have positive invader growth rates (eqn. 4 > 0), meaning that both competitors can invade when their competitor is at its equilibrium abundance, coexistence is predicted because both are buffered from extinction when at low abundances (Chesson 2000). We note that eqn. 4 can be modified to include a seedbank, as in eqn. 1.

To explore how interannual variation in germination phenology alters coexistence outcomes, it is necessary to know the distribution of phenological differences that species experience through time. Because this distribution is not known for any system that we are aware of, we simulated a simple scenario for which each species germinated in advance of the other by n days half of the time. For example, *V. microstachys* would germinate 5 days in advance of *V. octoflora* half of the time and 5 days behind half of the time, or 10 days in advance and 10 days behind, and so on. We then solved the mean of the invader growth rate (eqn. 4) for each species, or, equally, the long-term invader growth rate when the focal species is rare and the competing species is at equilibrium. This analysis is greatly simplified because the resident carrying capacity was unchanged by fluctuating germination dates, it was only interspecific competition that varied (see Results; Appendix S1). We supplemented this analysis by exploring how sensitive the outcome was to seed survival rate, which influences temporal coexistence (e.g., (Chesson and Huntly 1989; Abrams et al. 2013).

## Results

Each species produced a greater number of seeds per individual when it germinated earlier than the other species (Fig. 2; significant species × germination time(*F*_1,200_ = 629.9, *P <* 0.001)). *Vulpia microstachys* showed a two-fold increase in seed production when it germinated 10 days earlier vs. 10 days later than *V. octoflora*, whereas this difference was ten-fold for *V. octoflora* (Fig. 2). We found no effect of planting time on the number of seeds produced when each species was grown alone in monoculture (Fig. S2; non-significant time and time ×species (*P* > 0.2)), meaning that intraspecific competition was independent of planting date.

**Figure 2.**
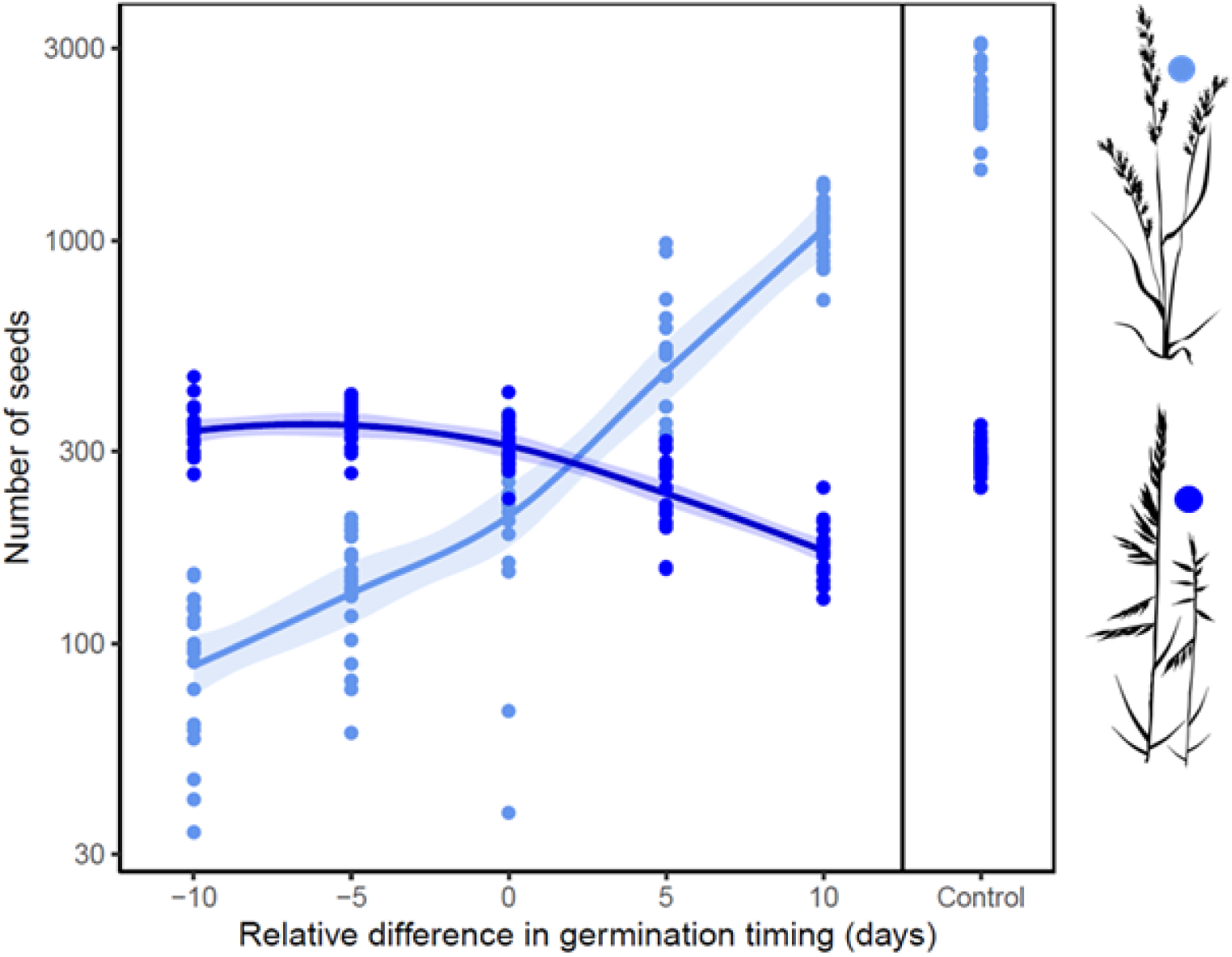
Differences in germination timing increase seed production for the earlier species. Plants produce two (*V. microstachys*, dark blue) to ten times (*V. octoflora*, light blue) the number of seeds when they germinate ten days in advance of the other species relative to when they germinate ten days after the other. Germination timing is negative when *V. microstachys*s germinates earlier. Each data point shows the number of seeds produced per experimental pot with 11 individuals of the focal species and its competitor. Control plots are monospecific.

Increases in seed production due to early germination were not counteracted by shifting seed mass. Although seed mass varied with differences in germination time, each species produced larger seeds on average when they germinated earlier (Fig. S3; significant germination time × species interaction (*F*_1,100_ = 25.8, *P* < 0.001)), reinforcing the seed number trends. Though present, seed mass changes were small relative to seed number trends, increasing 1.06-and 1.11-fold for *V. microstachys* and *V.octoflora*, respectively, between the earliest and latest germination times (Fig. S3). Because of this relatively small change in seed mass, and the unknown consequences for seed mass on competition, we did not incorporate seed mass into calculations of coexistence.

The effect of germination separation on niche differences and fitness differences was asymmetric (Fig. 3A)—both differences increased with germination separation when *V. microstachys* germinated first, but these differences first decreased and then increased when *V. octoflora* germinated first. This asymmetric effect can be understood by examining invader growth rates (Fig. 3B). Fitness differences were smallest when invader growth rates intersected, at approx. 4-5 days germination separation, and niche differences largely paralleled these fitness differences.

**Figure 3.**
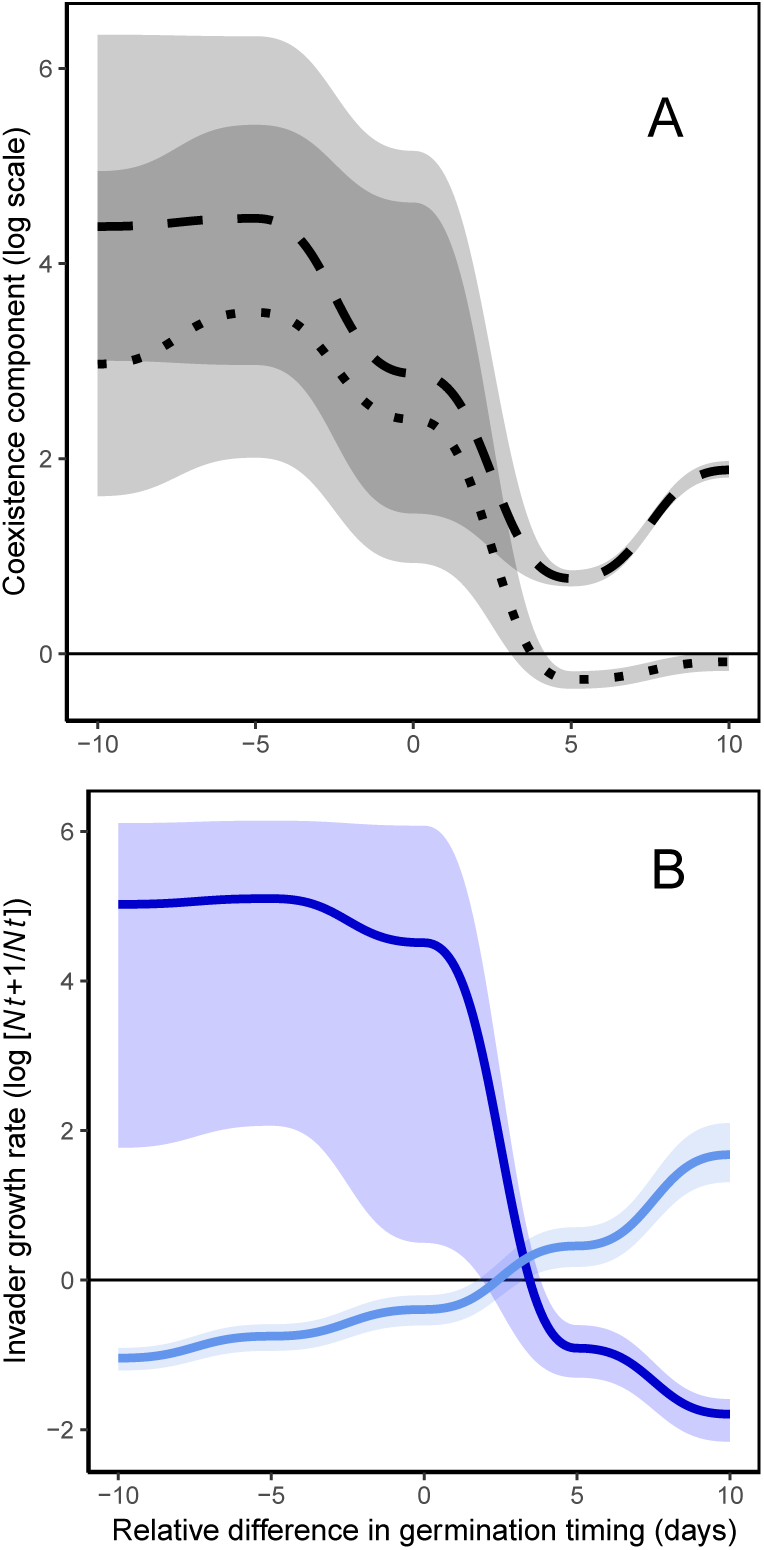
Differences in germination timing structure on fitness differences, niche differences, and invasion. (A) Fitness differences (dashed line; eqn. 3) and niche differences (dotted line; 1/*ρ* from eqn. 2) are greatest when phenological differences are large and cause *V. microstachys* to germinate first (left side of panel A). These components have the property that niche differences must exceed fitness differences for stable coexistence in a given environment. Minimum fitness differences coincide with maximum niche overlap, meaning that the components that limit coexistence are each favoured by different phenological pairings. (B) Greater phenological differences in germination increase low-density growth rates of the earlier germinating species, with *V. microstachys* shown in dark blue and *V. octoflora* in light blue. Coexistence appeared to be possible when shifts in germination timing caused a change in the superior competitor (approx. 4 day difference). In general, the negative correlation in low density growth rates between species shows that consistent differences in relative germination timing causes larger fitness differences than niche differences. Lines in both panels represent medians of data and envelopes delineate the 25th and 75th credible interval. Phenology effects were measured at 5 day intervals (−10, −5, 0, etc.) and lines between points were extrapolated using a weighted function.

The overall effect of differences in germination timing was that each species could invade and exclude the other when its germination was sufficiently in advance of the other (Fig. 3B; coexistence was only possible at 4 days). Interestingly, invader growth rate responses to germination timing were also qualitatively different for these species. *Vulpia* octoflora rates increased linearly as germination advanced, whereas for *V. microstachys*, invader growth rates were constant up until *V. microstachys* had a 5-day head start, and only showed small changes up until equal germination (0 day difference),upon which it decreased sharply (Fig. 3B). This nonlinear response was caused by the invasion growth rate of *V. microstachys* being limited by its maximum finite rate of increase, rather than by competition, when it germinated earlier than *V. octoflora*.

Our analysis of invader growth rates (eqn. 6) indicates that among year variation in interspecific competition via germination timing could promote long-term coexistence. Since differences in per capita interspecific competition (the only parameter in eqn. 1 to vary with germination separation) favoured different species in different scenarios, fluctuations through time might prevent exclusion of one species by the other from being realized. Mathematically, this occurs even in the absence of other fluctuations, because fluctuations in interspecific competition reduce its geometric mean (the second half of eqn. 4), and thereby increase the long-term invader growth rate (Appendix S1).To test this hypothesis, we simulated invasion potential of each species when germination timing fluctuated among years, giving one species an advantage only half of the time. We found that the weaker competitor on average, *V. octoflora*, could persist under the large fluctuations, where germination timing shifted between ± 7 or more days per year (Fig. 4). In contrast, *V. microstachys* persisted regardless of the amount of year-to-year variation. Thus, stable long-term coexistence was only predicted under scenarios of large fluctuations in relative germination timing (∼15 day or greater difference in relative timing from year to year).

**Figure 4.**
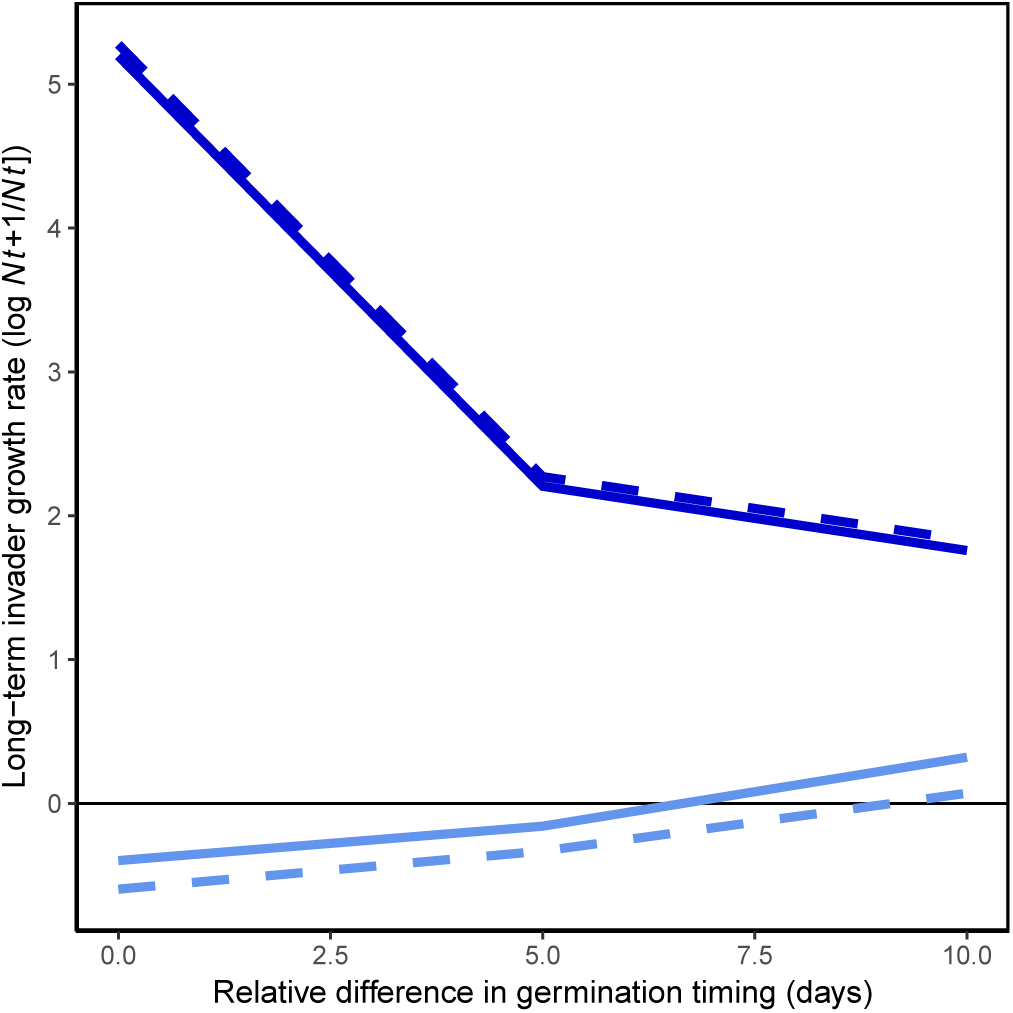
Large changes in the relative timing of germination among years favours coexistence. The invasion potential of both species when conditions cause one species to germinate earlier half the time and the other species to germinate earlier the rest of the time. The more extreme the differences in germination timing, the more coexistence is favoured; both species are predicted to persist if the annual differences in germination timing are seven or more days and all dormant seeds survive (solid lines), or greater than nine days even with no dormant seed survival (dashed line). Colours are light blue for *octoflora* and dark blue for *V. microstachys*. See methods for details.

## Discussion

Phenological differences between species have the potential to promote coexistence through increasing niche differences or preclude coexistence through increasing fitness differences. Thus, predicting how species’ phenological differences affect species coexistence is not clear, especially when those phenological differences fluctuate through time (Carter et al. 2018; Rudolf 2018; Satyanti et al. 2019). Our experiment shows that, although phenological differences in germination timing between two *Vulpia* species contribute to both niche and fitness differences, increases in fitness differences outweigh any increases in niche differences so long as phenological differences are consistent among yearsthe net effect is to limit coexistence. The phenologically-driven shift in fitness differences allowed the otherwise inferior competitor, *V. octoflora*, to invade and exclude *V. microstachys*. Indeed, our results suggest that long-term coexistence through phenological differences was only possible through fluctuations in phenological timing that effectively alternated the identity of the dominant competitor from year to year. Together, these results suggest that apparently contrasting research concluding that phenological differences limit coexistence (e.g. Godoy and Levine 2014) or promote coexistence (e.g. McKane et al. 1990) may be due in part to the temporal scales considered.

Our research clarifies how consistent phenological differences reduce coexistence by primarily affecting fitness asymmetries, with more dramatic phenological differences leading to larger fitness differences. This result echoes the conclusion of a recent field-based study on phenology that is, to our knowledge, the only other phenological research to disentangle coexistence mechanisms (Godoy and Levine 2014). Despite these apparent similarities between studies, phenological differences conferred fitness advantages in opposite scenarios—we found that earlier phenological timing caused species to be more competitive, whereas Godoy and Levine (2014) found the opposite. These opposing findings may be explained by two differences in how phenology was examined. First, in our study, we examined differences in phenology through separation of germination timing, whereas Godoy and Levine (2014) contrasted species that flower and reproduce at different times of the year (early spring vs. summer) despite germinating concurrently. In other words, our experiment provided earlier species with a ‘head start’ on resource uptake and competition (Ross and Harper 1972), whereas in their experiment, species with later phenology had a demographic advantage due to an extended growing season which allowed for resource uptake after the early species had senesced. Second, our experimental approach isolated the impacts of germination timing, whereas their approach captured phenological differences that correspond with suites of traits that influence reproductive rates (e.g. rooting depth and stem height) (Godoy and Levine 2014). As a result of these differences between studies, it is unclear if our common response of fitness differences to phenology reflects a broad pattern, or was simply coincidental, warranting future manipulative experiments in other species.

In natural conditions, fine scale phenological differences in germination can vary from year to year because germination is often tightly correlated to climatic conditions (Young et al. 1981; Günster 1994; De Luis et al. 2008; Levine et al. 2011). Nonetheless, germination cues that appear particularly effective for some species do not always coincide with optimal climate during the entire growing season, particularly when conditions suitable for germination are decoupled from those suitable to growth (Donohue et al. 2010). For example, Kimball et al. (2010) found that although the Sonoran Desert is becoming warmer and drier through time, species composition is shifting in favour of species that germinate and grow under colder conditions. The timing of winter rains which initiate germination have shifted to later in the year (December vs. October) causing plants to germinate under colder conditions even if average annual temperatures are on the rise. Our study suggests that, even when germination timing is independent of environmental conditions, early germination can provide a sufficient advantage to alter competitive outcomes.

We additionally found that the two *Vulpia* species differed in their sensitivity to separation of germination timing, illustrating that even closely related species respond differently to small changes in germination phenology by accruing different absolute advantages with early germination. The competitively dominant species *V. microstachys* had large impacts on *V. octoflora* in all treatments, reducing seed production compared to the monoculture control treatment even when germinating ten days after *V. octoflora* (Fig. 2). By contrast, *V. octoflora* showed minimal impact on *V. microstachys* seed production compared to the monoculture control treatment when *V. microstachys* germinated first (Fig. 2, 3). These differences suggest that greater phenological separation would disproportionately benefit the weaker competitor, as only*V. octoflora* has potential to further increase its low density growth rate when phenological separation increases beyond the range manipulated in our experiment. These differences in competitive effect and maximum seed production reflect differences in seed size and number; *V. octoflora* produces almost ten times as much seed as *V. microstachys* in the absence of competition (Fig. 2), however *V. microstachys* seeds are approximately five times heavier. Overall, the asymmetry in the importance of phenological timing for our *Vulpia* species suggests that differences in species’ traits that determine coexistence outcomes, such as seed size and fecundity, are likely to also influence how phenology alters competitive interactions.

A fitness advantage conferred to a species by arriving in advance of others is commonly referred to as a ‘priority effect’ (Fukami 2015). Priority effects are well studied but tend to be measured over time scales that span generations, such as when a second species arrives after an earlier arriving species has reached its equilibrium density (Peay et al. 2012). Measuring priority effects over shorter timescales, within the time scale of a generation, is less common (but see Black and Wilkinson 1963; Cleland et al. 2015) despite this being a relevant timescale for priority in life history transitions to alter competitive outcomes. We show how differences in germination timing cause strong priority effects, reversing which species is competitively excluded with as little as a 5 day separation. While previous work has shown that competition may be altered by germination timing (e.g. Aarssen 1989; Bergelson and Perry 1989; Cleland et al. 2015), it is surprising that the effects we document are large enough to be comparable to those caused by separation over generations (Fukami 2015).

Unlike priority effects that are produced over longer timescales from positive-density dependent population growth, and never promote local coexistence (Ke and Letten 2018), we show that within-season priority through germination differences could actually promote long-term coexistence if germination hierarchies vary among years. Specifically, we found that fluctuating conditions that lead to each species germinating in advance of the other in different years can promote coexistence, even though coexistence is not possible if germination timing is consistent among years. Previous research suggests that early and late rains favour different plant species (Wainwright et al. 2011). While this hypothesis has not been tested explicitly, our results based on empirically-parameterized simulations, coupled with widespread differences in germination timing commonly found in nature, indicate that it is likely important for understanding species coexistence.

Finally, climate change is expected to shift the relative timing of species’ life history events, which viewed through the lens of our experimental results, could impact patterns of species coexistence, and as a corollary, species composition of ecological communities (Kimball et al. 2010; Levine et al. 2011). We show that increased differences in germination phenology are unlikely to confer niche differences that increase coexistence and diversity, instead increasing the likelihood of competitive exclusion by the early species. This finding offers a critical link to help predict the ecological consequences of observed phenological shifts among competitors that might arise due to climate change. At the same time, we also show how among-year variability in germination phenology could facilitate coexistence in the long term, a plausible outcome of climate change for some species given predicted increased in interannual climate variability (IPCC 2013). Our research highlights the need to distinguish between chronic shifts in relative timing of phenological events and fluctuations in relative timing that may produce qualitatively different outcomes for competing species. Greater resolution on how shifts in phenological traits alter coexistence across ecological communities, and the temporal scales over which these shifts are likely to have an impact, are an important next step for predicting local consequences of climate change.

## Acknowledgments

We thank B. Hall, A. Petri, and A. Rego for assistance in the greenhouse. Research funding was provided to BG by NSERC (DG 2016-05621) and personal funding to RMG by NSERC CGS-D, the Biodiversity Research Centre, and Killam Trust.

