## Supplemental Materials for "An experimental manipulation of species’ phenologies overturns competitive hierarchies"

**Appendix S1 - Effects of fluctuating germination**

In the simulations we report, we imagine that germination timing fluctuates between years so that one species germinates *x* days in advance of the other half of the time, and *x* days after the other for the other half. In this case, the mean low-density finite rate of increase for species *i* is simply the geometric mean of the low density finite rate of increase. Without a seedbank, this is given by:

$$\sqrt{\frac{g_{1i}\lambda_{1i}}{1+g_{1j}\alpha_{1ij}N_{1j}}\frac{g_{2i}\lambda_{2i}}{1+g_{2j}\alpha_{2ijp}N_{2j}}} \qquad \text{(S1)}$$

Where the number signifies whether species *i* germinates first (1) or after (2) species *j*, and $N_j$ gives the equilibrium abundance of species *j* when alone. When all parameters except interspecific competition are constant through time, eqn. S1 simplifies to:

$$\frac{g_i\lambda_i}{g_jN_j\sqrt{\left(\frac{1}{g_jN_j}+\bar{\alpha}_{ij}\right)^2 - Var\,(\alpha_{ij})}} \qquad \text{(S2)}$$

From equation S2, it is apparent that an increase in the temporal variance of interspecific competition reduces overall competition, all else being equal. In the case where $g_jN_j$ is large, this fluctuation in interspecific competition will cause the multi-year effect of interspecific competition to become negligible when the its temporal coefficient of variation approaches 1 (i.e., $Var(\alpha_{ij}) \sim \bar{\alpha}_{ij}^2$). Importantly, because the variance in interspecific competition lowers long-run competitive effects of the resident, increasing this variation through temporal fluctuations in germination phenology will increase the long-run, low density growth rate of both invaders if the mean competitive effects of the residents stay fairly constant.

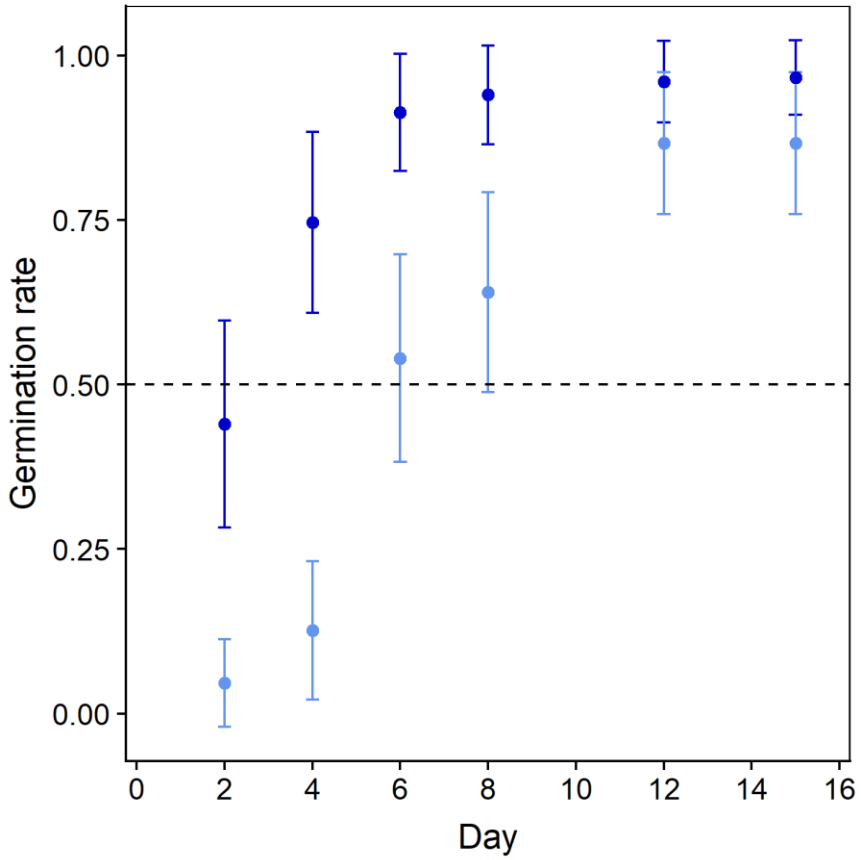


**Figure S1.** Germination timing of *V. microstachys* (dark blue) and *V. octoflora* (light blue). Error bars are ±standard deviation. The time to 50% germination (dashed black line) was approximately two days for *V. microstachys* and six days for *V. octoflora*. To control for any effect of taking early/late germinating seedlings, seedlings were transplanted on these days for each species.

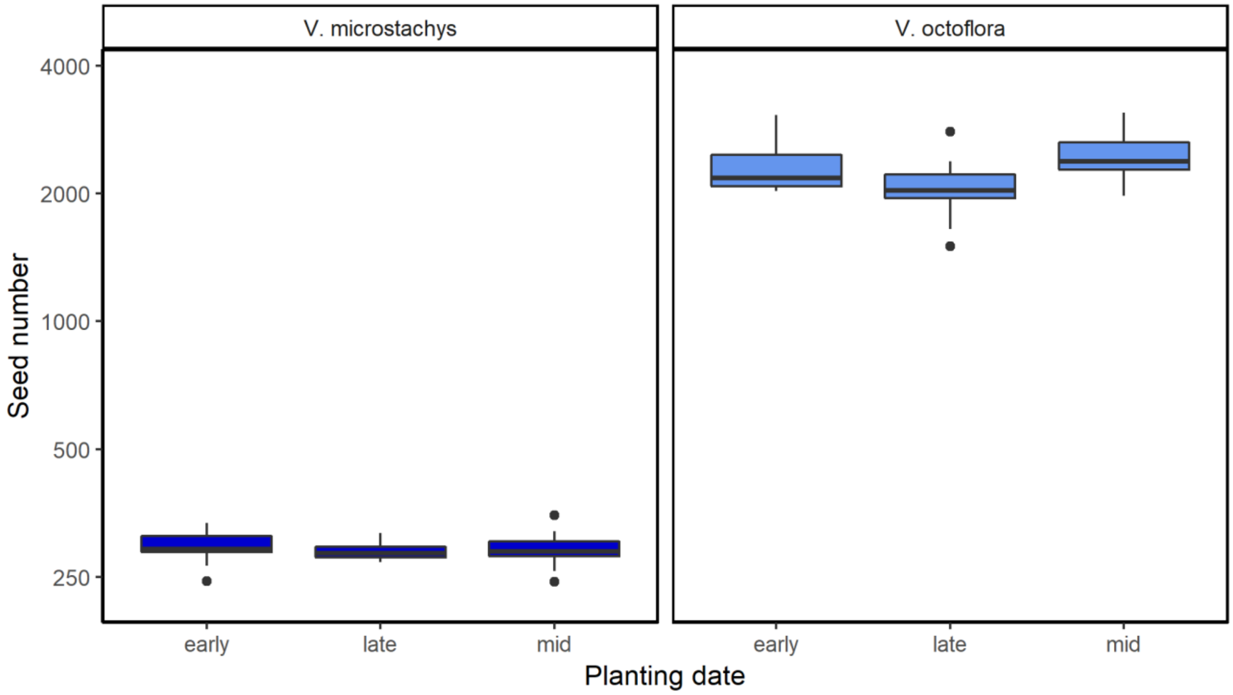


**Figure S2.** Pots in the monoculture control treatment showed no effect of planting date on seed production. *Vulpia microstachys* (left side) produced less seed on average when competing against conspecifics than *V. octoflora* (right side) but neither showed a trend with time ($P > 0.2$). Labels for germination timing coincide with whether controls were germinated to match the earliest treatments (early), at the same time as the midpoint treatments (mid), or for the treatment planted last (late).

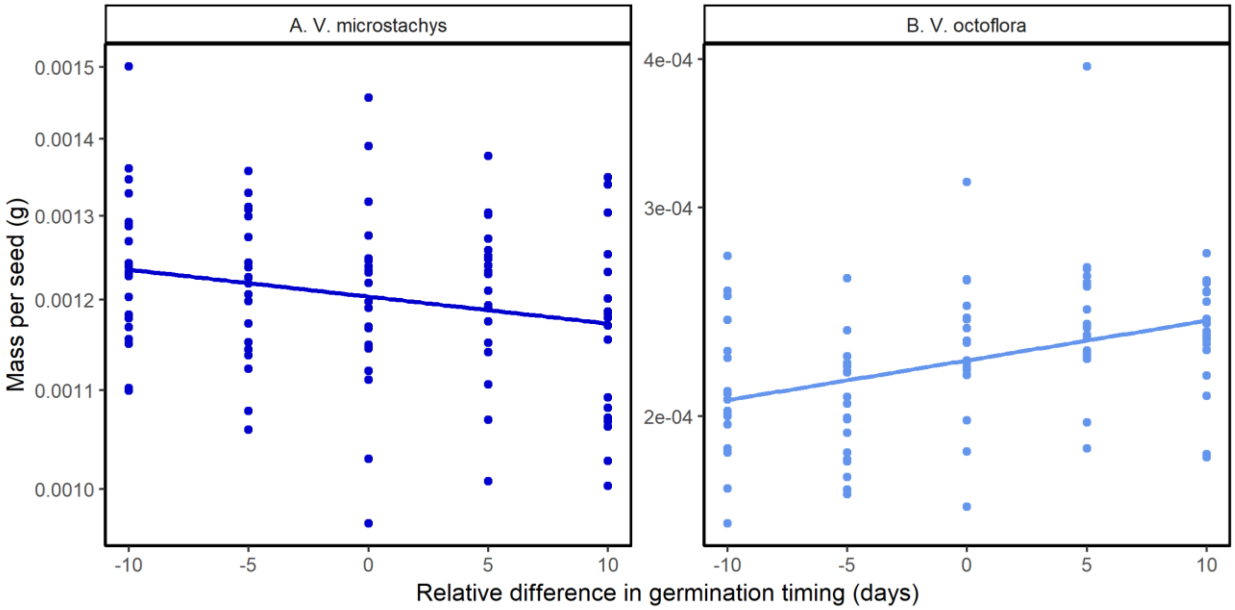


**Figure S3.** Differences in germination timing increase mass per seed for the earlier species. Mass per seed increases by 1.06 (A; *V. microstachys*) to 1.11 times (B; *V. octoflora*) when a species germinates ten days in advance of the other species relative to when it germinates ten days after the other. Note that the species differ in *y*-axis scales.
